# Neural correlates of individual facial recognition in a social wasp

**DOI:** 10.1101/2024.04.11.589095

**Authors:** Christopher M. Jernigan, Winrich A. Freiwald, Michael J. Sheehan

**Affiliations:** Laboratory for Animal Social Evolution and Recognition, Department of Neurobiology and Behavior, Cornell University; Ithaca, NY, 14853, USA; Laboratory of Neural Systems, The Rockefeller University, New York, NY 10065, USA

## Abstract

Individual recognition is critical for social behavior across species. Whether recognition is mediated by circuits specialized for social information processing has been a matter of debate. Here we examine the neurobiological underpinning of individual visual facial recognition in *Polistes fuscatus* paper wasps. Front-facing images of conspecific wasps broadly increase activity across many brain regions relative to other stimuli. Notably, we identify a localized subpopulation of neurons in the protocerebrum which show specialized selectivity for front-facing wasp images, which we term *wasp cells*. These *wasp cells* encode information regarding the facial patterns, with ensemble activity correlating with facial identity. *Wasp cells* are strikingly analogous to face cells in primates, indicating that specialized circuits are likely an adaptive feature of neural architecture to support visual recognition.

**One-Sentence Summary:** We identify a localized population of neurons specifically tuned to wasp faces in a social wasp that has independently evolved individual facial recognition analogous to the face cells of primates.

## Main Text

Adeptly navigating the social environment requires animals to modify their behavior depending on who they are interacting with. Given the importance of social interactions for fitness, there has been a longstanding interest in the evolution and mechanisms that mediate social cognition, especially recognition abilities (*1–8*). Social stimuli are often highly salient, with enhanced social discrimination or recognition abilities described in multiple species (*9–12*), and by specialized regions of the vertebrate brain (*13–16*). These observations have given rise to the idea of the ‘social brain’, circuits in the brain dedicated to processing social information (*17–19*). Some of the best evidence for socially selective neural circuits comes from the face-selective cell populations that mediate social recognition in primates (*16, 20–23*). Whether other taxa develop dedicated neural populations specialized for social recognition is not known but could provide evidence for their relevance in social recognition more broadly.

Like primates, female *Polistes fuscatus* paper wasps have evolved individual facial recognition that they use to mediate cooperation and conflict within their small societies (*24, 25*). Females in this species have individually distinctive facial color patterns that signal individual identity (Fig. 1A) (*24*). *P. fuscatus* paper wasps are specialized to discriminate faces above other visual objects (*10*), but this ability requires a holistic image of a paper wasp, i.e. behavioral discrimination is only achieved when faces are presented in the context of a whole wasp body (*26*). Remarkably, specialized face discrimination is not found in other closely related species of paper wasps that lack individual facial recognition, suggesting specialized processing of faces is a recent adaptation in *P. fuscatus* (*10, 27*).

**Fig. 1.**
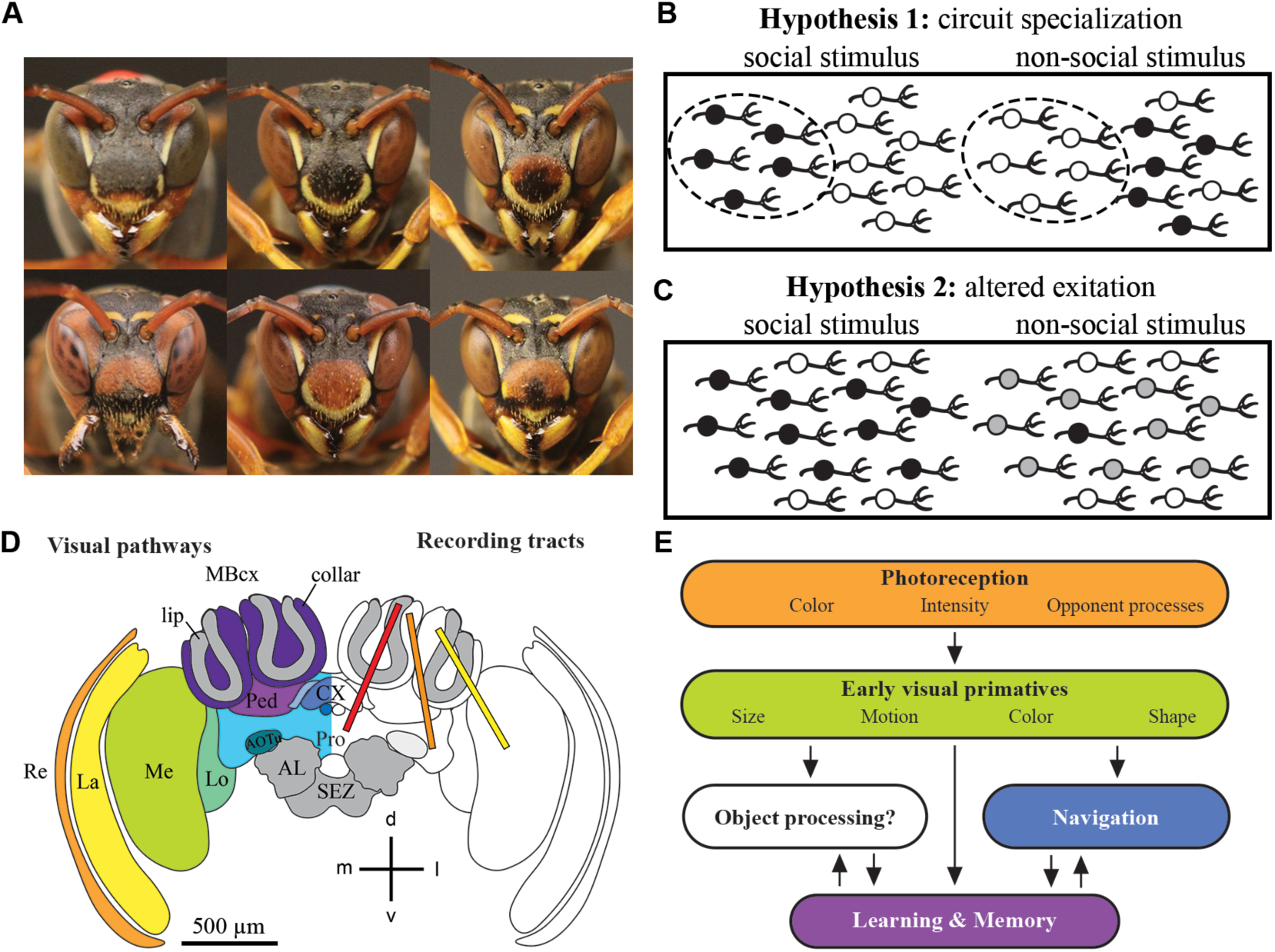
Proposed social selectivity outcomes among neurons in the brain and *Polistes fuscatus* as a model to explore social processing in the brain. (**A**) Example photos of unique facial patterns present on the faces of female *P. fuscatus*. (**B**) Diagram of circuit specialization hypothesis in which social and non-social stimuli are processed by different neural populations. Shading denotes high neural firing. Dashed ellipse denotes dedicated circuit. (**C**) Non-mutually exclusive hypothesis diagram in which social and non-social stimuli are both responded to above baseline, but neurons have altered excitation leading to selectivity for social over non-social stimuli. Shading denotes neural firing, black=strong neural firing, grey=moderate neural firing. (**D**) Schematic reconstruction of the wasp brain. *Left:* Colors denote visual neuropils, grey represents neuropils that canonically do not respond to visual information. Light blues denote central brain processes including the protocerebrum and optic glomeruli, dark blues denote the central complex, violets denote mushroom body processes, and warmer colors denote earlier optic lobe processes. *Right*: colored boxes denote the three recording tract locations explored. Lateral tract in yellow in the left hemisphere primarily covering optic lobe visual responses with some lateral mushroom body also recorded. Central tract in orange covering the central brain from the mushroom body calyx to optic lobe outputs to the lateral protocerebrum likely including optic glomeruli. Medial tract in red covering the central brain mushroom bodies, a small region of the medial protocerebrum, and input circuits to the central complex. Each wasp was recorded using only a single tract. Note Lateral and central tracts were recorded in the left hemisphere and medial tract was recorded in the right hemisphere. Below is a simplified schematic of visual processing in the insect brain with colors roughly matching where in the brain there is evidence of these computations occurring. Object processing remains unknown in the insect brain. Acronyms: Re= retina, La= lamina, Me= medulla, Lo= lobula, Pro= protocerebrum, AOTu= anterior optic tubercle, Ped= pedunculus of the mushroom body, MBcx= mushroom body calyx, lip= lip of the mushroom body calyx, collar= collar of the mushroom body calyx, CX= central complex, including the fan-shaped body, protocerebral bridge, and noduli. AL=antennal lobe, SEZ= suboesophageal zone, d=dorsal, v=ventral, m=medial, l=lateral. Scale bar denotes 500 μm.

The specialized cognitive responses to holistic wasp images strongly suggest that images of wasps generate distinct patterns of neural activity compared to other general image categories. One hypothesis is that facial discrimination in paper wasps is mediated by neural circuits that are specialized for images of wasps, similar to the face-cells of primates (*28*). If this is the case, we would expect to find visually responsive neurons in paper wasps that show distinct responses to front-facing images of wasps but are unresponsive or respond differently to other images (Fig. 1B). Based on behavioral data, hypothetical wasp specialized neurons should be sensitive to the shape and color of paper wasps (*10, 26, 29*). An alternative hypothesis is that cognitive specialization for facial recognition arises not because of specialized circuits but because social stimuli lead to increased excitement of generic circuits (*30, 31*). In this case, units that respond strongly to wasp images would also show elevated responses to other non-wasp image categories, albeit at a lower intensity (Fig. 1C). While these hypotheses are mutually exclusive for any given neuron, both types of neural firing patterns could exist among neurons within a brain, i.e., an overall increased responsiveness to social stimuli and a subset of neurons that show more extreme and specialized responses akin to specialized face patches in primates. Thus, using an independent evolution of vision and facial recognition compared to primates, we looked for object processing-like circuits with a focus on socially selective responses throughout the visual pathway of *P. fuscatus* wasps (Fig. 1D).

### Abundant neural responses to front-facing wasp images in the brain of *P. fuscatus*

To identify how wasp brains respond to images of conspecifics, we conducted large scale multi- channel extracellular electrophysiological recordings using 64-channel silicon probes (Cambridge Neurotech). We recorded a total of 794 visually responsive units across three recording tracts in 18 female *P. fuscatus*: a lateral tract primarily in the medulla and lobula of the optic lobe; a central track including the mushroom bodies, lateral protocerebrum, and optic glomeruli which receive inputs from the optic lobe; and a medial tract targeting mushroom bodies, central complex, and medial protocerebrum (Fig. 1D). We presented a stimulus set containing a broad range of natural images, geometric shapes, and images of wasps and their silhouettes to head-fixed paper wasps (Fig. 2A). Each stimulus category contained multiple unique images presented with a standardized minor looming motion that mimics the natural behavior of wasps during social interactions (Fig. 2B, see Methods for additional details).

**Fig. 2.**
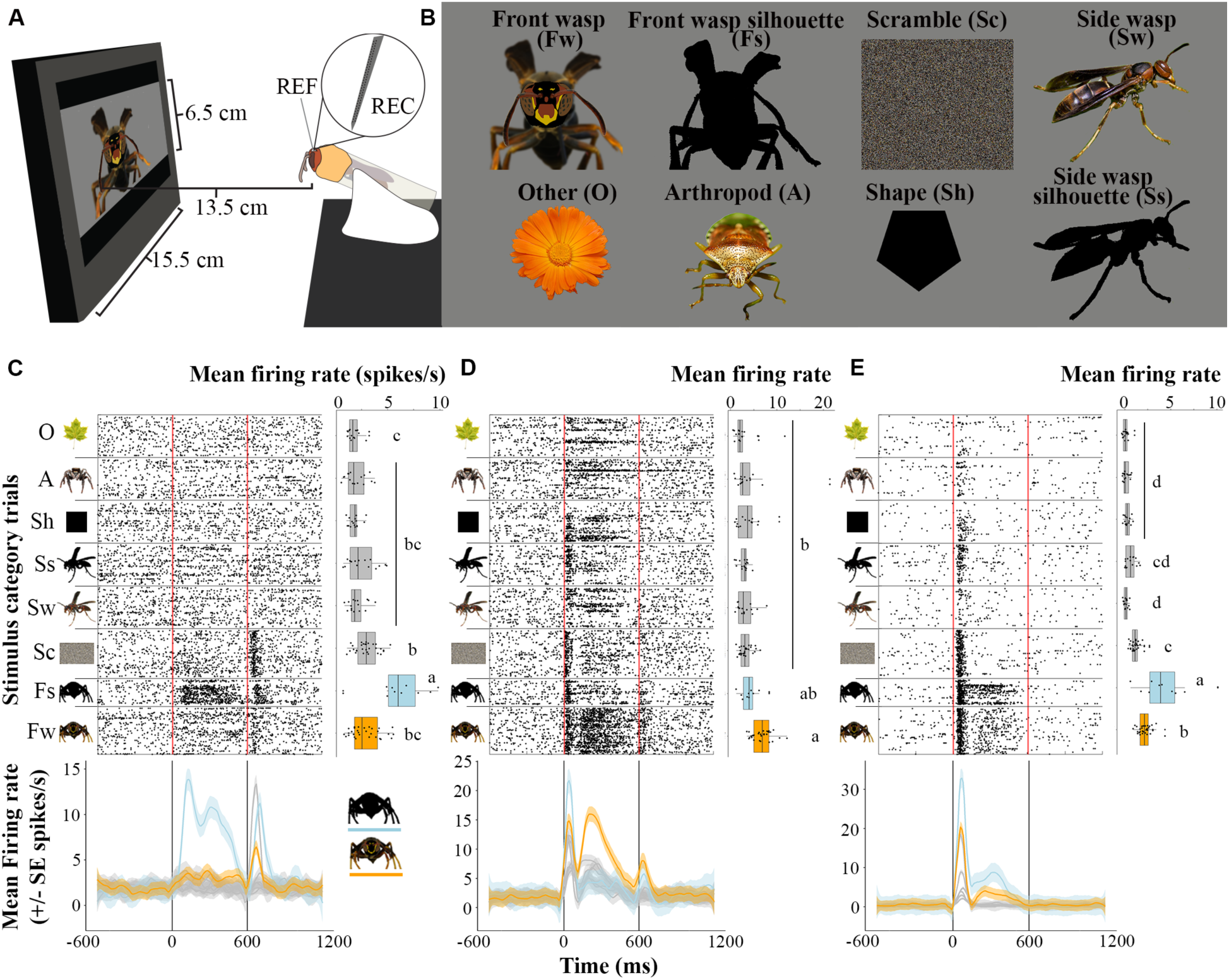
Neural responses to front-facing wasp shapes in the wasp brain. (**A**) Schematic of electrophysiological and stimulus display setup. (**B**) Example images of 8 stimulus categories used in stimulus set 1. (**C**-**E**) Example raster plots above, to the right are mean firing rate averaged for each stimulus category and below are mean firing rate for each stimulus category across time of units shown stimulus set 1. Each row in the raster plot is a unique stimulus presentation showing 2040 of the 2296 total presentations, made up of 690 of the total 706 unique stimuli sorted by the 8 stimulus categories highlighted on the y-axis (omitted car stimulus category for space and was a non-responded to stimulus). Each dot in the raster represents a unique action potential for a single unit aligned to stimulus presentation. Stimuli were presented for 600 ms and the vertical red lines represent stimulus onset and offset at 0 and 600 ms respectively during which the stimuli loomed slightly (see Supplemental methods for additional details). To the right of the raster plots, mean firing rates for each unique image (front-facing wasp averaged by unique face) within each stimulus category for the unit in the raster plots to the left are shown. Stats represent ANOVA analysis of mean firing rate ∼ stimulus category and letters denote p<0.05 significant difference by Tukey HSD post hoc analysis. Below are stimulus-onset aligned mean firing rates with GAM smoothing across time for each image category shown above. Color denotes stimulus category for each unit’s response front-facing wasp is highlighted in orange and front-facing wasp silhouette in blue. (**C**) Example unit 219 from wasp 1, which predominantly responds to the front-facing wasp silhouette stimuli. (**D**) Example *wasp cell* unit 270 from wasp 2, which predominately responds to the front-facing wasp stimuli. (**E**) Example *wasp cell* unit 365 from wasp 3, which responds predominantly to both the front-facing wasp silhouette and the front-facing wasp stimuli. Acronyms: REF=reference wire, REC=multichannel electrophysiological silicon recording probe, O=other, A=non-wasp arthropod, Sh=shape, Ss=side wasp silhouette, Sw=side wasp, Sc=scramble, Fs=front-facing wasp silhouette, Fw=front-facing wasp.

We reliably found visually selective cells that were highly responsive to front-facing wasp images, including both the silhouette and fully-colored wasp stimuli, across subjects and recording tracts (Fig. 1 C-E, figs. S1 & S2). Strong responses to front-facing wasp images or silhouettes would also be expected if the units are broadly tuned to large shapes or wasp colors. However, highly wasp-responsive units did not respond as strongly to wasp-colored geometric shapes or even wasp images or their silhouettes viewed from the side, suggesting that these cells are selective for the salient forward orientation of wasps (Fig. 1 C-E, figs. S1 & S2). Further these units were not responsive to just any complex image with visual heterogeneity, i.e., other front-facing arthropods nor other objects (Fig. 1C-E, figs. S1 & S2). Taken together, the firing patterns indicate the presence of units selectively responding to socially relevant stimuli in a view-dependent manner. Many units that were especially responsive to wasps showed responses to a range of stimuli at the onset of stimulus presentation but showed distinct persistent firing dynamics specific to front-facing wasp images (Fig. 2 DE, fig. S1BC). We consistently found strong responses toward front-facing wasp images across multiple animals and additional stimulus sets (fig. S2), indicating that strong responses to front-facing wasp images are a robust and frequently detected feature of visually responsive neurons in paper wasps (found in every recording with >10 visually responsive units).

### Generalized increased firing in response to socially salient stimuli across the brain

To assess the overall extent of specialization across all visual units, we calculated a selectivity index that captures the extent to which units have highly selective responses to a particular image category. Using this metric, positive scores for an image category indicates that on average a given unit fires to those images more than the average of all other image categories. A score ≥0.33 indicates that a unit fired at least twice as much to a category compared to the background average, indicating strong selectivity (*32, 33*). We examined the distribution of selectivity for the 794 visually responsive units we recorded using a single stimulus set.

Front-facing wasp stimuli matched stimuli that had been used to behaviorally assess discrimination in *P. fuscatus* previously (*10, 29*) and were therefore our target stimuli, and silhouettes of those front-facing wasp stimuli acted as shape controls that lacked wasp color patterning. Scramble stimuli had been designed to disentangle front-facing wasp shape and color patterning responses. Since scrambles share color content with front-view wasp images without containing any shape information, joint response preferences to the two stimulus categories would reveal color selectivity. Joint response preferences to front-facing wasp and silhouette stimuli, but not scrambles, would reveal invariant front-facing wasp shape selectivity. Joint responses to side-views of wasps and front-facing views of wasps these would have been view invariant responses to conspecific images. Selective responses to the two silhouettes and the geometric shapes would reveal high contrast shape or edge detectors. Finally, strong non-selective responses to all image categories would simply reveal looming detectors.

We found most image category selectivity distributions were centered at or near zero and only rarely had evidence of selective responses (2%, 18/794), indicating that visual regions of the wasp brain are neither broadly excited nor inhibited by these categories ruling out much of the response types listed above such as high contrast shape or edge detectors or widespread loom detectors (Fig. 3A, table S1). We do see a slight shift in selectivity for side-view silhouette images of wasps with most (9/12) also being selective to front-view wasps, suggesting there may be some, albeit rare, view-invariant wasp shape selectivity in our dataset (Fig. 3A, table S1). However, three image categories exhibited a marked rightward shift, with their median selectivity substantially greater than zero: the front-facing wasp, its silhouette, and scrambled- versions of the front-facing wasp images (Fig. 3A, table S1). All three elevated-response stimulus categories had evidence of many modestly excited units that were not necessarily specialized but tended to show increased responses to social stimuli (29% or 228 that had between 1.25x and 2x selectivity to front-facing wasp, 40% or 314 to front-facing silhouette, and 32% or 255 to scramble). We found similarly elevated responses to these three image classes with other stimulus sets (table S2) and with other statistical approaches to categorize elevated firing patterns (table S3). Together this suggests we find broadly elevated responses to front-facing conspecific shapes as well as wasp color selectivity across the visually responsive units in our dataset.

**Fig. 3.**
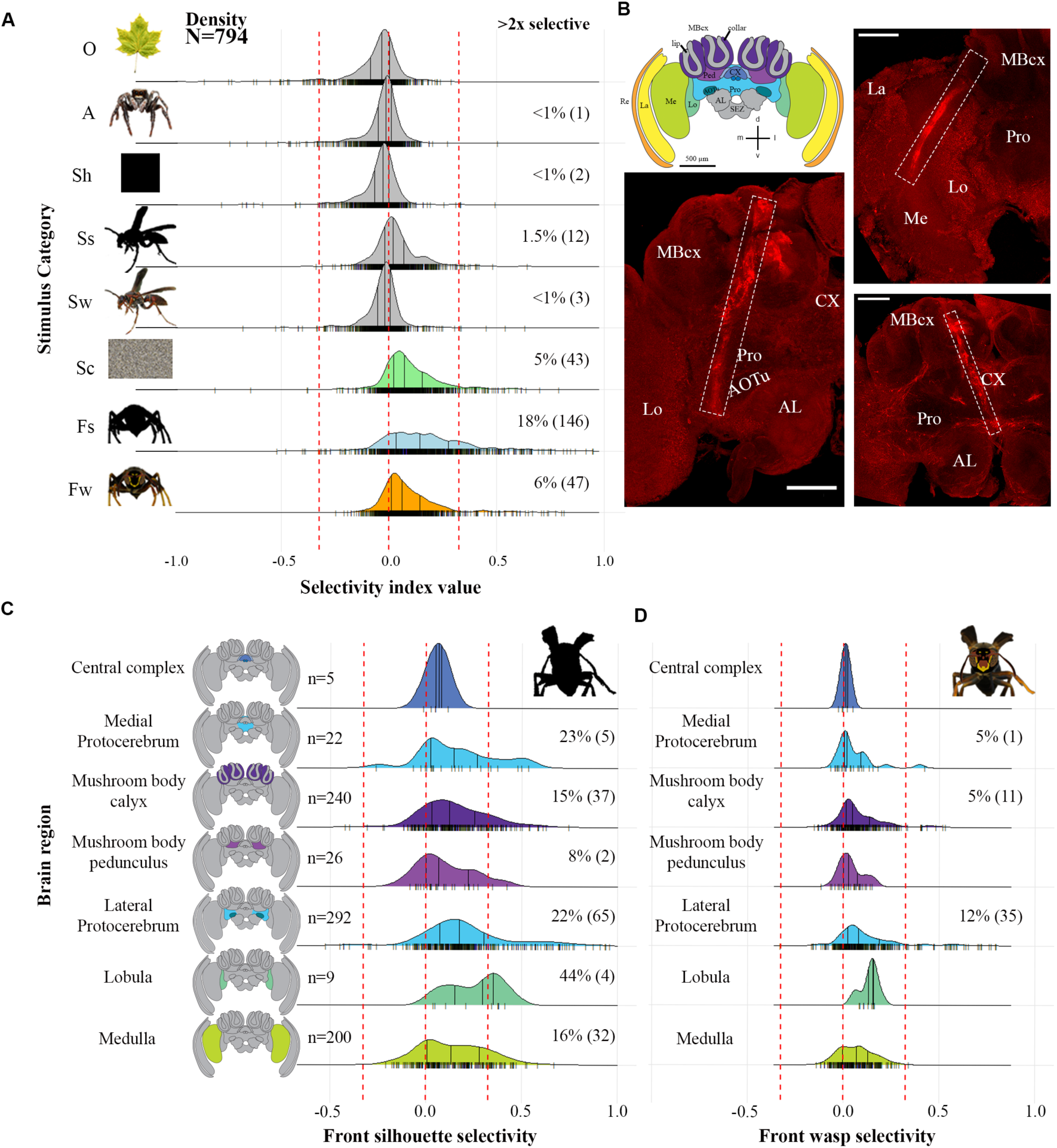
Broad selectivity to front-facing wasp shapes across the wasp brain with selectivity to front-facing wasp stimuli in the lateral protocerebrum. (**A**) The ridge-line plot shows the distribution of stimulus category selectivity for each of the 794 visually responsive units recorded using stimulus set 1. Three highlighted categories have distributions substantially shifted away from zero, indicating that they generally drive increased responses across the wasp brain. See methods for additional information regarding selectivity index calculation. Numbers to the right denote percentages (and number of units in parentheses) of units that show >=2x selectivity. (**B**) Schematic of the wasp brain as well as example fluorescently labeled recording tracts for the central tract (bottom left), lateral tract (upper right), and medial tract (lower right). Dashed boxes denote staining tracts. Scale bars denotes 200 μm. (**C**) Distribution of selectivity to front-facing wasp silhouettes across regions of the wasp brain. (**D**) Distribution of selectivity to front-facing wasp images across regions of the wasp brain. Numbers to the right in C and D denote the total number of visual units record in each neuropil and percentages (and number of units in parentheses) to the right denote the percentage of units in each neuropil that show >=2x selectivity to front-facing silhouette or wasp stimuli in C and D respectively. Acronyms: Re=retina, La=lamina, Me=medulla, Lo=lobula, Pro=protocerebrum, MBcx= mushroom body calyx, AOTu= anterior optic tubercle, AL=antennal lobe, CX=central complex, SEZ=suboesophageal zone.

### Wasp selective units are localized to the lateral protocerebrum

We next examined the selectivity of units by their estimated neuropil location in our three recording tracts (Fig. 3B). First, across all visual units in our dataset we see broadly elevated selectivity to the front-facing wasp silhouettes across the entire wasp brain (Fig. 3C, ∼20% of all visual units across all brain regions). This suggests that in the case of these paper wasps, it may be said that much of the brain is broadly responsive to front-facing wasp shape, supporting in part the non-specialized social processing hypothesis. Similar broad projections of courtship associated circuits have been described in the fruit fly (*34, 35*), suggesting this may be a common feature across insect brains.

Next, we looked at responses selective to the front-facing wasp stimuli themselves. Most units that showed strong selectivity to colored images of front-facing wasps were also selective to the front-facing wasp silhouette (98%, 46/47, table S1). These front-facing wasp selective units were found almost exclusively in the lateral protocerebrum or mushroom bodies (Fig. 3D) with only one other unit being found in the medial protocerebrum where we rarely found units that responded to any of the presented stimuli. Of all the front-facing wasp selective units, 75% (35/47) were located in the lateral protocerebrum near optic glomeruli and these patterns are consistent across additional experiments and stimulus sets tested as well as analyses based on relative channel position rather than neuropil (fig. S3).

Together these finding suggests an abundance of highly selective units to both front-facing wasp shape and wasp color localized in the wasp lateral protocerebrum, near the approximate depth of the anterior optic tubercle, the other optic glomeruli, and their outputs. One of these optic glomeruli, the anterior optic tubercle (AOTu), has previously been shown to have increased volume relative to mushroom body visual calyces in socially reared animals compared to those that lack visual social experience in this species (*36*). In other insects the AOTu is involved in color processing (*37*), female-female aggression (*38*), and projects to higher order sensory integration centers (*39*). Additionally, we see a similar pattern of an abundance of scramble image selective units in the lateral protocerebrum (fig. S4), which further suggests this region may be involved in color pattern processing. We focus our further analyses on this group of localized highly wasp-selective units located in the lateral protocerebrum, hereafter referred to as *wasp cells*.

### *Wasp cells* respond to wasp shape and facial pattern variation

These *wasp cells* are clearly responding to the shape of a front-facing conspecific both with and without wasp color patterning. However, they appear to respond to both the shape and color independently, why do some *wasp cells* (37%, 13/35, table S1) respond selectively to both the scramble images that have wasp color but lack the wasp shape as well as the silhouettes that have the wasp shape but lack the wasp color? These two stimuli seem as if they cannot be more different.

When we look at selectivity over smaller bin windows instead of summed over the entire response window (0 to 1080 ms) we see selectivity to scramble images differs significantly from those to the front-facing wasp and its silhouette particularly after the initial onset response (Fig. 4A-D, fig. S5, GLM: selectivity ∼ stimulus * time). This is particularly pronounced in those units with persistent firing patterns (Fig. 4B&D) as well as both the most and least scramble selective *wasp cell* (fig. S5). This suggests that these units are generally tuned to wasp colors, however when the wasp shape is not present they quickly stop responding.

**Fig. 4.**
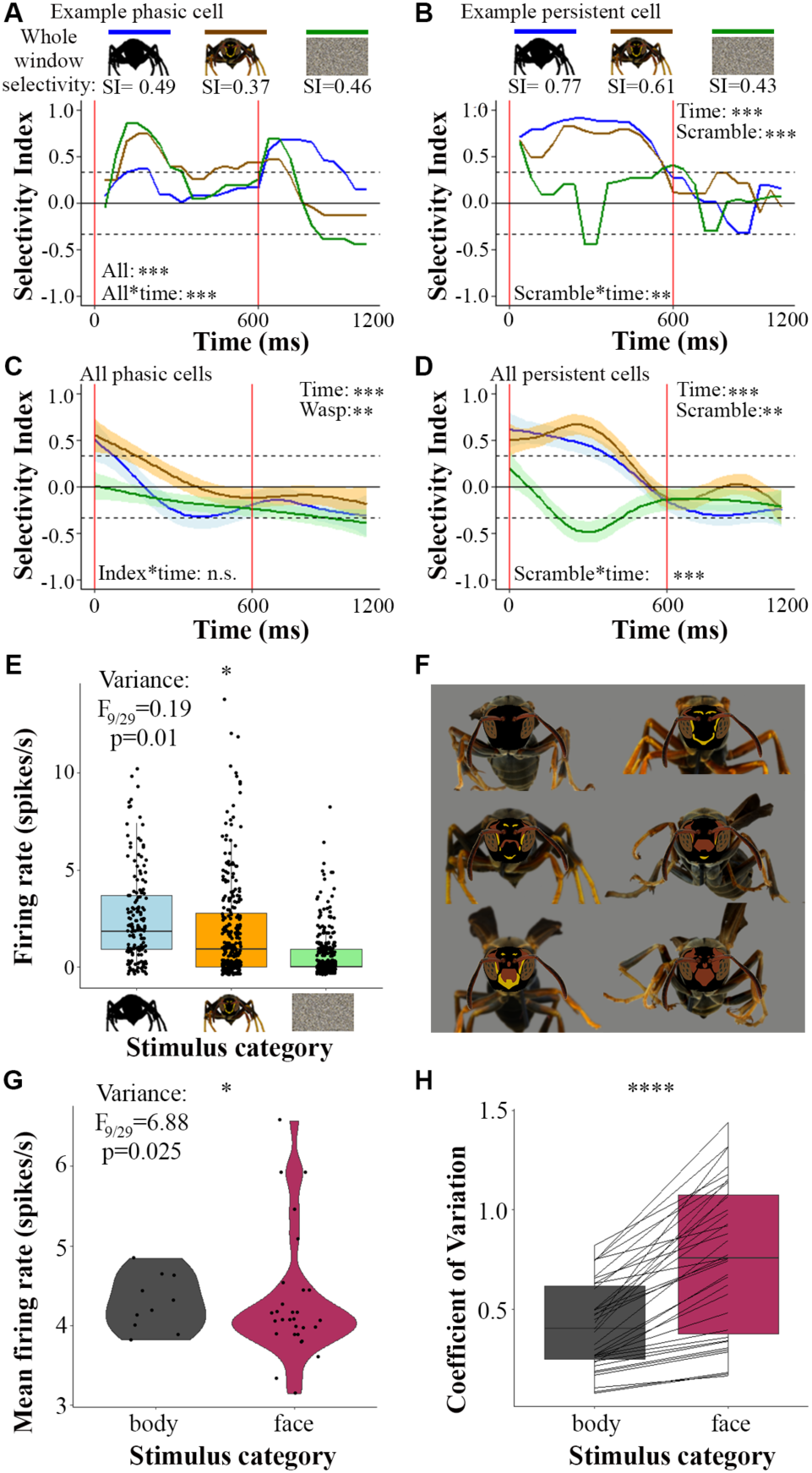
*Wasp cell* responses to front-facing wasp shape and color stimuli. (**A**) Example phasic spiking *wasp cell* 186 from wasp 4 whose raster is plotted in Fig. S1A and its selectivity in 40ms bin windows over the whole stimulus response window (0-1080 ms) to front-facing silhouette (blue), wasp (orange), and scramble (green). Selectivity index (SI) values for this cell’s whole response window are displayed above. Stats denote significance running GLM analysis, see supplementary methods for additional details. (**B**) Example persistent *wasp cell* 365 from wasp 3 whose raster is plotted in Fig. 2E and its selectivity in 40 ms bin windows as in A. Line plots in A and B are smoothed using a median selecting a value across 3 bins. (**C**) Model fit of selectivity to over time for the 18 phasic *wasp cells* front-facing silhouette (blue), wasp (orange), and scramble (green). Stats denote significance running GLM analysis, see supplementary methods for additional details. (**D**) Model fit of selectivity to over time for the 17 persistent spiking *wasp cells* as in C. (**E**) Example *wasp cell* unit 330 from wasp 3 and its per-trial firing rate across the stimulus presentation window from 0-1080 ms for all presentations of front-facing wasp silhouette, front-facing wasp, and scramble image stimuli. Stats denote F-test comparing variance across average peak firing rate stimulus categories. (**F**) Example subset of 6 unique bodies each with a unique wasp face of the total 30 faces and 10 bodies within stimulus set 1. See fig. S4 for full set of bodies and faces. (**G**) Example *wasp cell,* unit 206 from wasp 4, and its mean firing rate to the 10 presented bodies and 30 presented faces. Stats denote F-test comparing variance across face responses and body responses. (**H**) The firing rate coefficient of variation across 10 presented bodies versus the 30 presented faces for each of the 35 *wasp cells* shown stimulus set 1. Lines connect the same unit in each bar plot, line color denotes relationship: black=greater variation to face than body. Stats denote paired t-test of coefficient of variation across all *wasp cells*. Colors denote stimulus category: blue= front-facing wasp silhouette, orange= front-facing wasp, green= scramble, grey= front-facing wasp body, maroon= front- facing wasp face.

What about responses to silhouettes? *Wasp cells* are highly selective to the shape of a front-facing wasp, however somewhat counter intuitively their average firing rate is higher to silhouettes, which lack color pattern information and contain only the wasp shape. However upon further examination of the range of firing to wasp images and silhouettes we find that while *wasp cells* tend to fire more on average to silhouettes, most (85.7%, 30/35) show the strongest response to a normal wasp image (Fig. 4E). Overall, 27 *wasp cells* (77%, 12 of 35 significantly so based on a paired F-test) show a broader range of peak firing in response to color-patterned wasp images compared to the silhouettes of the same image, suggesting that these units receive and process color information. The broader overall range of firing by most neurons in the presence of color suggests that the natural range of wasp color patterns have the potential to both increase and suppress firing relative to a large black silhouette of the wasp. An alternative possibility is that the silhouette may be a super-stimulus, as insect vision has been reported to be particularly attuned to high contrast dark shapes (*40–43*) like the silhouettes shown here.

Given the overall responses in the wasp brain to wasp shapes and colors, we hypothesized that *wasp cells* might be tuned to body shape (posture) and/or facial patterning (color) and thus show variable firing rates across one or both of these components in stimulus images. If *wasp cells* responded more variably to different bodies, it would suggest possible postural signal encoding, which is known in this species (*44*). Alternatively, if *wasp cells* varied more across the different facial patterns, it would suggest selectivity for the smaller area of the wasp image that contains the relevant facial identity information. To test the role of body shape versus facial coloration, we presented 30 faces on 10 different bodies for a total of 300 unique combinations (Fig. 4F, fig. S4AB). We find most units showing significantly greater variation to faces than bodies (54%, 19/35, paired F-test) (example cell: Fig. 4G, table S4). Though notably the remaining non-significant *wasp cells* all still have greater variation in firing rates to facial patterns than to body shape in absolute terms, this is exemplified by looking at all 35 *wasp cells* as a population, which shows significantly increased variation in response across faces compared to body shape (Fig. 4H, paired t-test, t=4.681, df=54.595, p=1.93x10^-5^). The wasp face covers a relatively smaller portion of the total image (23% of the image on average, fig. S6A), yet more variation in the responses is driven by the color patterns on the faces compared to the posture of the body. These results indicate that the firing dynamics of *wasp cells* are particularly sensitive to facial pattern variation.

### Population correlates of facial identity discrimination by *wasp cells*

Increased variation in responses to facial patterning rather than body posture suggests that *wasp cells* might encode information about facial identity. In primates, individual facial identity is encoded by populations of face cells, with different neurons having distinct tuning properties and preferences of different aspects of facial variation (*45–47*). As a result, face cell populations can accurately differentiate among many unique faces in a multi-dimensional face space (*45*). Examining the preferences of individual *wasp cells*, we find that the peak firing rate significantly differs among faces across units (ANOVA, peak firing rate ∼ face + body, face p<0.05), and that face-preference ranking appears to be idiosyncratic and highly inconsistent across units even within the same animal (fig. S6C). This result indicates that units may have unique preferences for holistic faces or specific facial features. To further explore what aspects of facial features wasp cells may be attending to, we designed a new stimulus set which included 42 unique combinations made up of variations in 3 standardized facial pattern elements and tested this on further *wasp cells* identified in the same region of the wasp lateral protocerebrum (Fig. 5A). We again found *wasp cells* with peak firing that differs across holistic front-facing wasp images and that these units have idiosyncratic preferences across faces with highly preferred faces and highly unpreferred faces (Fig. 5B-M, z-score firing differs significantly from 0, unpaired t-test, p<0.05, fig. S7), with unique facial preferences among recorded *wasp cells* from the same animal (Fig. 5B,F). Furthermore, a subset of *wasp cells* in these recordings preferentially discriminate among single facial features, such as markings on the clypeus or the ‘eyebrow’ like stripe above the antenna (Fig. 5B-I). This additional stimulus set emphasizes variation in a small number of facial features, so apparent tuning preferences for individual features as opposed to more complex axes of facial variation may be a consequence of the stimulus set. However these patterns suggest that some *wasp cells* are tuned to particular axes of facial space as has been previously demonstrated in primate face cells (*45*).

**Fig. 5.**
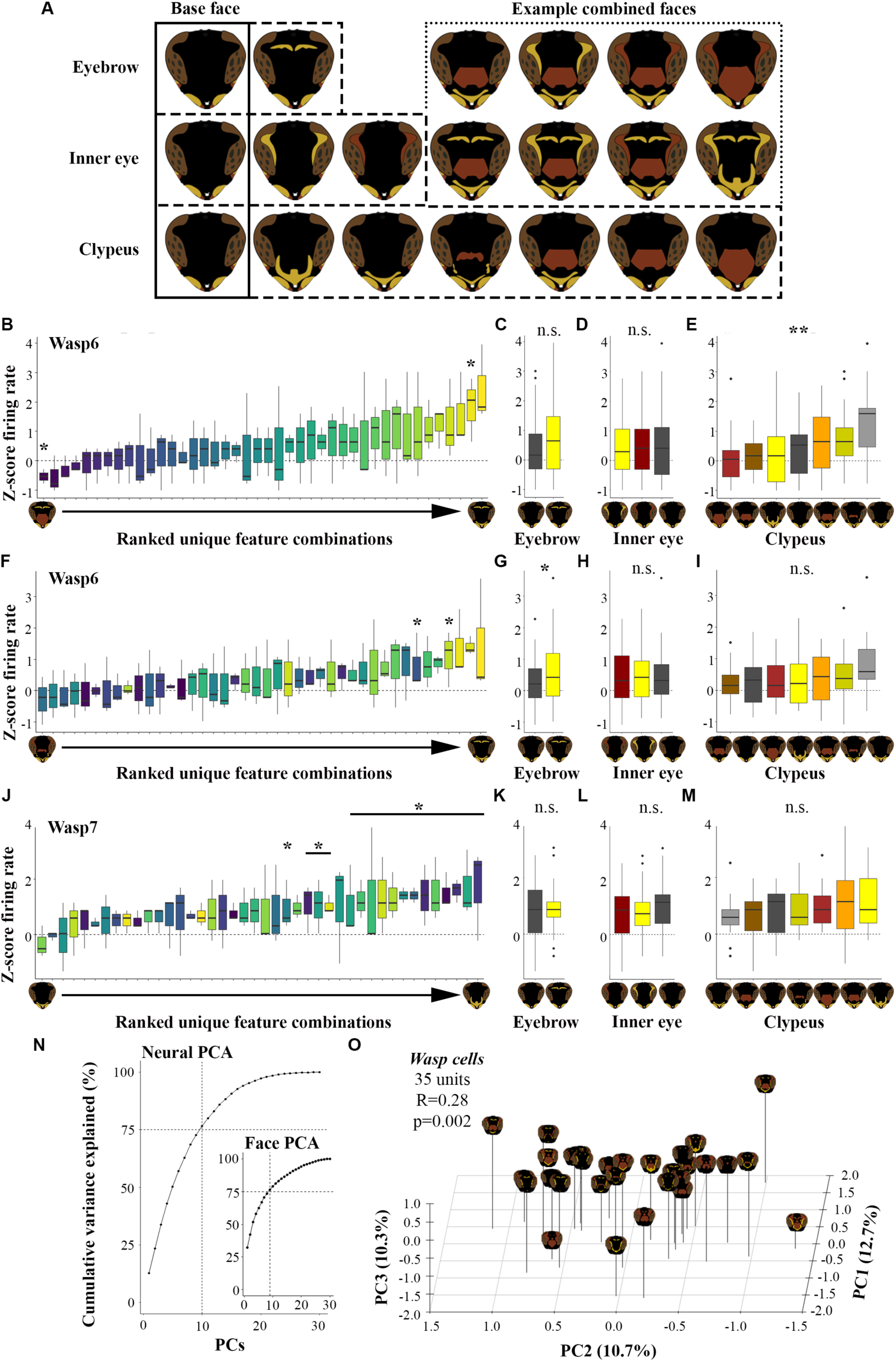
Unique idiosyncratic face and facial feature preferences of *wasp cells* and their phenotypic separation of presented wasp faces. (**A**) Parameterized face components used to make front-facing wasp stimuli in stimulus set 3. Eyebrow absent or yellow. Inner eye yellow, red, or absent. Six clypeal patterns or absent. Solid box outlines base face on which each facial region pattern was added. Dashed boxes outline each facial sub regions which we varied. Dotted line outlines example combinatorial faces showing one clypeal pattern and the 6 combinations with eyebrows and inner eye components as well as 2 additional unique combinatorial faces. These 2 “eyebrow”, 3 inner eye, and 7 clypeal patterns allowed the generation of 42 unique holistic facial patterns. (**B-E**) Z-score normalized peak firing rate for unique faces or facial feature shown to *wasp cell* unit 365 from wasp 6 shown stimulus set 3. (**F-I**) Z-score normalized peak firing rate for unique faces or facial feature shown to *wasp cell* unit 254 from wasp 6 shown stimulus set 3. (**J-M**) Z-score normalized peak firing rate for unique faces or facial feature shown to *wasp cell* unit 51 from wasp 7 shown stimulus set 3. (**B, F, J**) Plot of ranked z-score peak firing across unique holistic face combinations. (**C, G, K**) Plot of ranked z-score peak firing across unique eyebrow patterns. (**D, H, L**) Plot of ranked z-score peak firing across unique inner eye patterns. (**E, I, M**) Plot of ranked z-score peak firing across unique clypeal patterns. Stats for B, F, and J denote significant difference of z-score normalized firing rate for each unique facial combination from zero via unpaired t-test. Stats for C-E, G-I, K-M denote significant difference by ANOVA, model= Average peak firing rate ∼ eyebrow*inner eye*clypeus + body, for each facial component. (**N**) Plot of the cumulative variance explained for each principal component (PC) of the principal component analysis (PCA) of the 35 *wasp cells’* z-score peak firing to the 30 wasp faces shown in stimulus set 1. Line denotes 10 PCs are required to explain 75% of the total variance present across *wasp cells*. Inset plot shows the same analysis for wasp phenotype PCA (see methods for additional details) with 9 PCs required to explain >75% of the total variance in presented facial stimuli. (**O**) Plot of face locations in the first 3 PCs of the neural PCA for the 35 *wasp cells’* z-score peak firing to the 30 wasp faces shown in stimulus set 1. Stats denote significant correlation of facial distances across neural firing PCA space and facial phenotype PCA space via Mantel test with 100,000 permutations.

If single cells are tuned to particular axes of facial space, population ensemble firing patterns may also encode variation in the facial features to allow for individual recognition as has been described in primates (*45–47*). To explore this we ran a principal component analysis (PCA) of the mean peak firing rates to each facial stimulus for all *wasp cells* and we also created a multi- dimensional wasp face space using our stimuli (*25*). We found the neural principal component (PC) space to be highly multidimensional, taking 10 PCs to explain >75% of the variance in population stimulus responses from 35 *wasp cells* (Fig. 5N). Notably, the dimensionality of neural firing among the *wasp cells* closely resembles the dimensionality of phenotypic face space of presented stimuli, where 9 PCs are needed to explain >75% of the variation (Fig. 5N). If *wasp cell* populations can encode facial identity, we should observe a correlation between inter-face distances in phenotypic face space and the neural firing space of *wasp cells*. Remarkably, we find the distances between faces in neural and face PC space are significantly correlated (Fig. 5O, Mantel test, R=0.28, p=0.002, 100,000 permutations), indicating that *wasp cell* populations phenotypically separate faces similarly to our facial pattern analyses. Further, if we re-run this analysis including only *wasp cells* from the same animal (2 separate animals for which we have enough *wasp cells*) this significant relationship also holds independently for each population of *wasp cells* (fig. S8, Wasp3: 27 units, R=0.18, p=0.036, Mantel test, 100,000 permutations; Wasp4: 7 units, R=0.22, p=0.017). Further, the correlation between neural and face PC spaces only holds when considering responses to front-facing wasp images, and not when substituting the responses to scramble controls for which shape and pattern information is absent, but color proportions remain (Mantel test, p>0.42). This indicates selected *wasp cells* meaningfully differentiate among facial patterns *per se* and not simply color proportions in each image.

Next, we asked if the ability to discriminate among wasp faces wasp specific to the identified population we termed *wasp cells*? Using front-facing wasp selective units located in the mushroom bodies, rather than the lateral protocerebrum, we compared face distance relationships among neural and facial PC space, and found that these mushroom body units had no relationship with facial phenotype distance (11 units, Mantel test, R=0.08, p=0.187), nor did a larger number of mushroom body units using less restrictive selectivity threshold (0.24 versus 0.33, 26 units, Mantel test, R=0.08, p=0.184). Together these analyses indicate that facial pattern tuning is a distinct property of the *wasp cells* we have identified in the paper wasp lateral protocerebrum. These findings suggest that *wasp cells* are specialized and are performing similar, at the gross level, visual object computations to the fusiform face area of the primate brain. We hypothesize that this rather complex behavior could be achieved with a relatively small number of *wasp cells*, on the order of hundreds, given that with a relatively sparse number of *wasp cells* we can meaningfully separate distinct facial patterns. This fits with our working hypotheses that an optic glomerulus, such as the AOTu, could have been adapted to fit this role *in Polistes fuscatus* (*36*).

## Discussion

The *wasp cells* we have described here are strikingly analogous to primate face cells despite the vast evolutionary distance between primate and wasp lineages indicating that specialized neural populations are likely an adaptive and efficient architecture for visual identity processing. Paper wasp and primate lineages convergently evolved brains, eyes, sociality, and individual facial recognition. In both taxa, faces contain signals of individual identity (*24, 48–50*) and visually responsive neurons in these systems are selective for front-facing orientations of conspecifics that display identity signals (Fig. 2, figs. S1 & S2) (*23*). Further we can localize these *wasp cells* to the lateral protocerebrum (Fig. 3), near the optic glomeruli we had previously shown to be plastic to social experience (*36*), and that front-facing wasp selective units in the mushroom bodies do not show the same relationship in firing to displayed facial patterns. Like ensemble encoding of identity by primate face cells (*45*), we also find evidence that population firing patterns of *wasp cells* contain information on facial identity (Fig. 5, fig. S8). Additionally, among these we find some persistently firing units that show differential firing patterns to social and non-social stimuli (front-facing wasp shape vs. scrambles, shape, etc., Figs. 2E, 4A-D, figs. S1 B&C, S5) as has also been recently reported in primate face cells (*51*). Convergent evolution of similarities in specialized neural populations for visual identification between wasps and primates argues that specialized circuits are an adaptation for social recognition, even though developmental experiences influence face specialization in both taxa (*52, 53*). Studies probing the features of *wasp cells* and face cells will further clarify which aspects of recognition circuits may be lineage-specific versus shared optimal features of adaptive circuits in both taxa.

In addition to specialized circuits, social stimuli broadly drove excitation across visual brain regions in the wasp. These finding provide new insight into the debate about the role of specialized or dedicated circuitry for social behavior, which has involved diverse techniques at different scales across species There is growing evidence of social stimuli activating broad circuits across animal systems (*34, 35, 54–58*). In primates there is also evidence for face selectivity as early as V4 in the primate brain (*59*) and possibly earlier (*60*). There are two possibilities that could cause this pattern, there exists a broad feedback circuit that projects to earlier visual brain regions possibly driving this global enhancement and at least one cell type that fits this function is known to exist in the fruit fly brain (*35*). Alternatively, there may be neurons intrinsic to early visual centers that can detect and shift global responsivity when front-facing wasp images are being presented thus priming selectivity in later circuits. Regardless of how this global shift in responsivity is achieved, this work builds on a growing number of findings across animals that the social brain may be considered both a composite of specialized nodes and a modulator of whole brain state activity.

## Materials and Methods

### Animal housing and care

*Polistes fuscatus* wasp were wild caught from Ithaca, NY, USA. Wasps were provided *ad libitum* sugar and water, and housed under a 12/12 light/dark cycle. They also received blue, green, and violet colored paper for visual enrichment.

### Stimulus generation

Digital images of 16 natural objects including flowers, wasp nests, and barns were obtained from Wikimedia commons and personal photographs. Digital images of 16 cars, 16 front-facing views of non-wasp arthropods, and 16 side-views of *Polistes* wasps were obtained from Wikimedia commons. These images had their backgrounds manually removed in Adobe Photoshop (CC21), centered, and exported as a PNG to 504 by 324 pixel size at 72 dpi with a 50% grey background.

#### Stimulus set 1 wasp images

We created 15 unique front-facing *Polistes fuscatus* face patterns in Adobe illustrator (CC21) using photos of *P. fuscatus* faces. An additional 14 face patterns, plus a no-pattern face, were manually created using mixed features of other faces to cover a broader range of phenotypic *P. fuscatus* face space based upon analysis of 267 *P. fuscatus* faces (*25*). Wasp face patterns were manually placed in the correct position on a silhouetted *P. fuscatus* head manually created in Adobe Photoshop. All wasp pigments in face patterns were standardized to be identical colors in Adobe photoshop to standardize color and contrast on each face. Notably, *Polistes fuscatus* wasps do not contain any non-visible (ultraviolet) facial signaling components (*25*). Additionally, an identical standardized eye pattern and image of *P. fuscatus* antenna were also placed in identical locations for all front-facing wasp images. Photos were taken of 10 front-facing views of wasp bodies with heads removed. Backgrounds of each body photo were manually removed in Adobe Photoshop and photos were resized such that the thorax of each body was the same diameter and centrally placed. Each wasp face pattern and body combination was output with identical face, antenna, eye, thorax positioning. Each image was exported as a PNG to 504 by 324 pixel size at 72 dpi and placed on a 50% grey background.

Each body (excluding portions covered by face and antenna) and face pattern were independently analyzed in ImageJ for percent and number of pixels for each of the three wasp pigments: yellow, red-brown, and black. Custom Code was written in MATLAB to then randomly generate a random pixel image rectangles that covered the same maximal area as each body and contained the same pixel resolution and number of yellow, red-brown, and black pixels as each front-facing wasp view image.

Silhouettes for each body and each wasp side-view were created using the fill function using 100% black in Adobe Photoshop. Additionally geometric shape images were created in Adobe Illustrator and moved to Adobe Photoshop to resize and color to match the three primary wasp pigment colors used in facial patterns: yellow, red-brown, and black.

#### Stimulus set 2 wasp images

We photographed 12 unique wasp facial patterns with antennae removed from wasp collected in the Ithaca area. We then created cartoonized facial pattern masks in adobe illustrator for the patterning between the eyes including the eyebrows (frons), inner eye, and clypeus with each pattern color matching precisely the color in the photograph. We also created one standard cartoonized eye mask that was used with all wasp pattern stimuli. We then photographed 3 front-facing wasp bodies with heads removed and one wasp with antenna attached. From these images we created 4 image categories: front-facing wasp photo (photo of wasp face, antenna, and body), front-facing wasp pattern (cartoonized facial pattern mask on a head silhouette background, standard cartoonized eye, and antenna and body photos).

For this stimulus set each face cartoonized pattern/photo was only presented on a single body such that we cannot disambiguate body selectivity from face selectivity in this dataset. For this reason, we did not further analyze face pattern preference for this dataset. However, we can robustly identify wasp selective units as well as compare if these units discriminate stimuli using this dataset. We used this stimulus set to compare responses among photos, patterns, and silhouettes. Again, random pixel scramble images match the maximal height and width of the front-facing wasp bodies as well as the number of pixels of each color in the front-facing wasp images

Each body (excluding portions covered by face and antenna) and face pattern were independently analyzed in ImageJ for percent and number of pixels for each of the three wasp pigments: yellow, red-brown, and black. Custom Code was again applied to randomly generate a random pixel image rectangle that covered the same maximal area as each body and contained the same pixel resolution and number of yellow, red-brown, and black pixels as each front-facing wasp image.

#### Stimulus set 3 wasp images

We generated unique facial feature components consisting of 2 eyebrow regions (or frons: yellow vs. absent), 3 inner eye patterns that varied by color (yellow, red, or absent), and 7 unique clypeal patterns (6 unique patterns found in nature and one entirely black clypeus). Each of these features was combinatorially added on to a default base head with standardized antenna and mandibles to create 42 unique holistic wasp faces. Each of these 42 faces was then placed on 3 unique wasp bodies used in previous recording experiments.

### Stimulus Presentation

All above stimuli were saved as PNGs and then saved into a video file using custom code and the Psychtoolbox (*61, 62*) in MATLAB (*63*). Images were presented for 600 ms (36 frames) with 1200 ms between each image presentation. Images were maximally 504 by 324 pixels (6.5 by 15.5 cm) in size and centered at the 307 by 300 pixel position within a 1024 by 600 pixel, 7in monitor screen. Images loomed beginning at 0.8415 times the image size (424 x 273 px) and looming to a maximal size of the image size (504 x 324 px) at frame 28 in a sinusoidal pattern at a rate of *abs*(*sin*(0.4π\**t* + 1), where *t* is time in frames (60 fps). Video was generated, saved, and played to match the output of the stimulus monitor at a rate of 60 frames per second.

#### Stimulus set 1

For front-facing wasp images each wasp face and body combination was presented once per recording, leading to each face pattern being seen 10 times and each body being seen 30 times for a total of 300 unique front-facing wasp presentations. Each of these 300 front-facing wasp presentations also had a corresponding color and size controlled random pixel scramble image. All other object stimuli were presented 16 times. This gave rise to a total of 256 stimulus presentations (16 unique stimuli each presented 16 times) for each of the following stimulus categories: cars, natural objects, geometric shapes, front-facing non-wasp arthropods, side view *Polistes* wasps, and side view silhouettes of *Polistes* wasps.

This gives rise to a total of 2,296 stimulus presentations made up of 706 unique stimuli (600 of which are front-facing wasps and random scramble controls) in a 1 hr and 17 min long stimulus video with an additional 168 canonical stimuli not discussed. In a few recordings there were additional supplemental stimuli (not discussed) this went up to 2,872 stimulus presentations made up of 742 unique stimuli in a 1 hr and 31 min long video with additional supplemental stimuli. The order of each stimulus presentation was randomized for each animal.

#### Stimulus set 2

We created 8 primary stimulus categories: front-facing wasp silhouette (N=3), front-facing wasp pattern (N=12), front-facing wasp photo (N=12), Side wasp (N=6), Side wasp silhouette (N=6), cars (N=5), geometric shapes (N= 18), and random pixel scramble (N=12). All object stimuli were presented 16 times. In this stimulus set we presented 1,184 stimulus presentations made up of 74 unique stimuli in a 55 min long stimulus video with additional stimuli not discussed. The order of each stimulus presentation was randomized for each animal.

#### Stimulus set 3

The stimulus set consisted of combinatorial front-facing wasp images (N=126, 42 faces on 3 bodies), front-facing wasp silhouette (N=3), side wasp images (N=8) and side wasp silhouettes (N=8) arthropod (N=8), arthropod silhouette (N=8), geometric shapes (N=8), scramble images (N=3), other natural objects (N=8), cars (N=8), and human faces (N=8) were shown. All object stimuli were presented 10 times along with 168 canonical stimuli not discussed and additional *Polistes* wasp species stimuli also not discussed. We only discuss *wasp units’* responses to front-facing wasp images in this manuscript, but used comparable stimulus categories to stimulus set 1 (did not include arthropod silhouette or human faces) to identify *wasp cells* via front-facing wasp selectivity >0.33 and depth >400 μm on the recording shank. Order of each stimulus presentation was randomized for each animal.

### Electrophysiological procedure

Animals were chilled at 4°C for ∼5 min, this did not cause full anesthesia and wasps never stopped moving but were slowed. After the mild cold anesthesia, wasps were placed within a small cut tube with their head and the top of their thorax placed above the upper edge of the tube. The animal’s body and head was fixed in place using pliable sticky wax (Wipmix, No. 10847). A small incision was cut in the back of the head capsule using a breakable blade creating a small window between the eye and the ocelli, exposing the dorsal surface of the mushroom bodies. The trachea and neural sheath were removed in a small area to allow for neural probe insertion.

Cambridge neurotech H5 and H8 neural probes were used (Cambridge Neurotech silicon neural probes, ASSY-77) with a silver reference wire. Recording electrode shanks were labeled with micro-ruby dye (Invitrogen, D7162) for localizing recording track. Electrode shanks were inserted through the mushroom bodies and aiming for 1 of 3 tracts (Fig. 1D). Recording sites varied slightly (∼50 μm) between animals, however all recordings followed the same general position and were confirmed via fluorescent dye imaging via confocal microscopy (Zeiss i710). After electrode shank insertion, a reference electrode was placed nearby within head capsule hemolymph above the tracheal sheath. Next a small amount of ringer’s solution (∼2-4 μL) was added in 15 min increments over the next 30-50 mins prior to recording to allow neural tissue to recover following electrode insertion. Animals were placed ∼13.5 cm away and directly facing the center of a small computer monitor (Loncevon, MODNXA6XQJVXHL, 7 in size, 1024*600 pixel resolution monitor, 60 Hz refresh rate). Immediately following this rest period, animals acclimated to the dark for ∼10 mins. Following this dark acclimation recording began. See table S5 for additional metadata information for each wasp and recording session.

### Electrophysiological acquisition system and spike sorting

Multichannel extracellular electrophysiological data was recorded across 64 channels, and two analog input channels, two stimulus alignment channels, at a rate of 25 kHz using an Intan systems acquisition system (intan technologies, LLC, RHD2000). Recording channels had 60 Hz noise filter and notch filters applied, additionally a white noise filter was applied during spike sorting.

Stimulus presentation timing was aligned via an audio signal (not audibly presented) that occurred during each stimulus presentation via an input to an analog channel in an acquisition system, additionally we had a backup alignment channel connected to a photodiode that allowed for the assessment of the beginning and end of stimulus video. This allowed for an independent way to identify the precise onset and timing of stimulus presentations for the entire length of the electrophysiological recording.

Data was saved in 1 min files and then combined into a single .bin file using Intan provided and custom code in Matlab. Spikes were then extracted into distinct clusters using *Kilosort3* (*64*) using default settings. All clusters or units were then manually checked using *Phy* (*65*). During the manual unit check, units were merged if: spike wave forms were highly similar, spike amplitude was within 20 μV, if dominant spike channel was the same or on a neighboring channel and the unit clearly jumped channels, and if unit spike timing was physiologically possible to be one neuron. Next all extracted spikes for each unit had their times aligned to each stimulus presentation using custom code in Matlab. For all units, raster plots and peristimulus time histograms (PSTH) were created. These plots were then manually checked to select units whose firing pattern changed in response to presented visual stimuli. These remaining visually responsive units were then selected for further analyses. Additionally, units were removed from further analysis if baseline firing was too high (mean inter spike interval was less than 4 ms), or the firing rate was too low to obviously discern response selectivity. Additionally, we selected a window of peak firing for each unit based upon PSTH plots.

### Data Analysis

We recorded a total of 794 visually responsive units across three tracts using stimulus set 1, and an additional 311 visually responsive units in the central tract using stimulus set 2 and 264 visually responsive units using stimulus set 3 across 28 wasps in total across all stimulus sets. All statistical tests were performed in R (*66*). ANOVA, t-tests, chi-square test, Pearson’s correlation tests, F-tests, and PCA analyses were performed using base R, and Mantel tests were performed in R using the ade4 package (*67*). Full models and comparisons are provided in main text. Additional data information and rate calculation information is provided below. We analyzed rates two ways, either via the total response window, including ‘on’ and ‘off’ responses, or via peak firing rate, selected for each unit in the ‘on’ window based on PSTHs. Generally, for inter-stimulus category measures we compared total response window rates (0 to 1080 ms) and for intra-stimulus category assessments, i.e. discriminating among images within a category, we only compared ‘on’ responses via peak firing rates (between 0 to 150 ms up to 0 to 600 ms—the full stimulus ‘on’ window—depending upon the firing dynamics of each neuron). Peak firing rate was also used to compare variation in firing across stimulus categories for identified *wasp cells*.

#### Stimulus category selectivity

We calculated binned total firing rate from 0 to 1080 ms for each visually responsive unit which included both ‘on’ and ‘off’ responses similar to Chang & Tsao (*45*). We then calculated a stimulus category selectivity index similar to Rolls (*32*) where for each stimulus category we calculated the mean total firing rate for each stimulus category vs. the other stimulus categories such that for each unit:

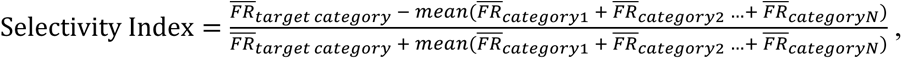

where *F̄R̄* is the mean firing rate and *N* is the number of stimulus categories. After initial discovery of bias for front-facing wasp, front-facing silhouette and scramble we excluded these responses from being directly contrasted against each other in this calculation and treated them as a single response category when treated as non-target in the index calculation, such that image categories were not over- or under-inflated due to this obvious bias across the nervous system. We defined selective units as those that had a selectivity index score >0.33, i.e. on average fired more than 2 times more to a target image category than the other image categories (*33*). We also calculated selectivity in 40 ms time bins from 0 to 1080 ms for *wasp cells*, and then performed a GLM analysis comparing selectivity across stimulus three stimulus categories: front-facing silhouette, front-facing wasp, and scramble. Generalized linear model (GLM) analyses in R (*66*) with stimulus category and time as interacting fixed effects. We also ran GLM analyses for each *wasp cell* as well as across *wasp cells* that were manually selected as either phasic firing or persistent firing *wasp cells,* based upon raster and PSTH plots. Additionally we compared selectivity relationships across units using Pearson correlations in R (*66*).

In addition to using the selectivity index, ANOVA tests were used to identify image selective units with p<0.001. We calculated binned firing rate during for the first and second 300 ms of each stimulus presentation for each visually responsive. We then ran an ANOVA (Firing rate ∼ stimulus category) for each unit such that a unit was identified as selective if the firing rate significantly greater for one category than the firing rate for all other image categories via Tukey HSD posthoc analyses. However, we found this approach to capture the same highly selective units but was less conservative than the selectivity index and therefore remained with units identified via selectivity index. Comparison of this approach versus the selectivity index approach is provided in table S3.

#### Wasp feature differences

For wasp feature analyses looking across body and face we only used *wasp cells* shown stimulus set 1 (N=35). To identify if units varied more in firing across faces than across bodies or vice versa we analyzed variance two ways. First, we performed an F-test for significant variance within each unit comparing variance across mean peak firing rate responses to each presented wasp face versus variance across mean response to each presented wasp body. Second, we compared coefficient of variation across mean firing rate across faces versus across bodies, such that each unit had a face coefficient of variation value and a body coefficient of variation value, we then compared these two coefficients via a paired t-test. Further we subsampled random pools of 10 faces going into the coefficient of variation calculation and across all subsamples also found highly significant differences in coefficient of variation with face responses having greater coefficient of variation than bodies across all runs, run 5 times.

To identify if units varied in firing across faces, we compared peak firing rate (window selected for each unit via PSTH plots) or full stimulus response window and used ANOVA to compare firing rates (firing rate ∼ face + body). Most units only differed in firing across the peak firing window. Additionally, we analyzed *wasp cells* recorded using stimulus set 3 (N=8) to identify facial feature discrimination using z-score peak firing rate and used ANOVA to compare firing rates across the interaction of each facial feature as well as body (z-score firing rate ∼ frons * inner eye * clypeus + body). Additionally, to compare if units varied in firing across holistic faces we used ANOVA as above (z-score firing rate ∼ face + body) as well as used an unpaired t-test to see if z-scored firing rate differed from zero, either significantly above or below, with Benjamini-Hochberg correction for multiple testing. These results indicated differential firing across faces.

#### Population firing rate comparison to phenotype space

To identify possible population level responses to presented wasp features we performed a principal component analysis (PCA) across all *wasp cells* and within single animals for which we had a sufficient number of recorded *wasp cells*. The input to the PCA analysis was the mean peak firing rate (peak window determined by PSTH) to each presented wasp faces. Thus, we obtained principal component (PC) values for each presented wasp face. Additionally, we used a Mantel test to correlate the distances of faces in neural PC firing space with their distances in phenotype space using the *ade4* package in R (*67*). We generated phenotype PCA space distances using the package *patternize* (*68*) in R following the procedures outlined in Tumulty et al. (*25*). We included two additional controls by repeating this analysis but used peak firing rate of *wasp cells* to scramble images as well as examining different population of wasp image selective units located in the mushroom bodies rather than the protocerebrum and repeated the above face neural PCA and phenotype PCA distance correlations utilizing these units to show that wasp selective units found in the protocerebrum are in fact unique.

## Acknowledgments

We thank Dr. James P. Tumulty for running phenotype distance scores for presented wasp face stimuli. We also thank Drs. Gaby Maimon, Nilay Yapici, Andrew H. Bass, and Jesse H. Goldberg for their thoughtful comments on early drafts of this manuscript.

## Author contributions

Conceptualization: CMJ, MJS, Methodology: CMJ, Investigation: CMJ, Visualization: CMJ, Funding acquisition: MJS, Project administration: CMJ, MJS, Supervision: MJS, WAF, Writing – original draft: CMJ, Writing – review & editing: CMJ, WAF, MJS

---

**Fig. S1.**
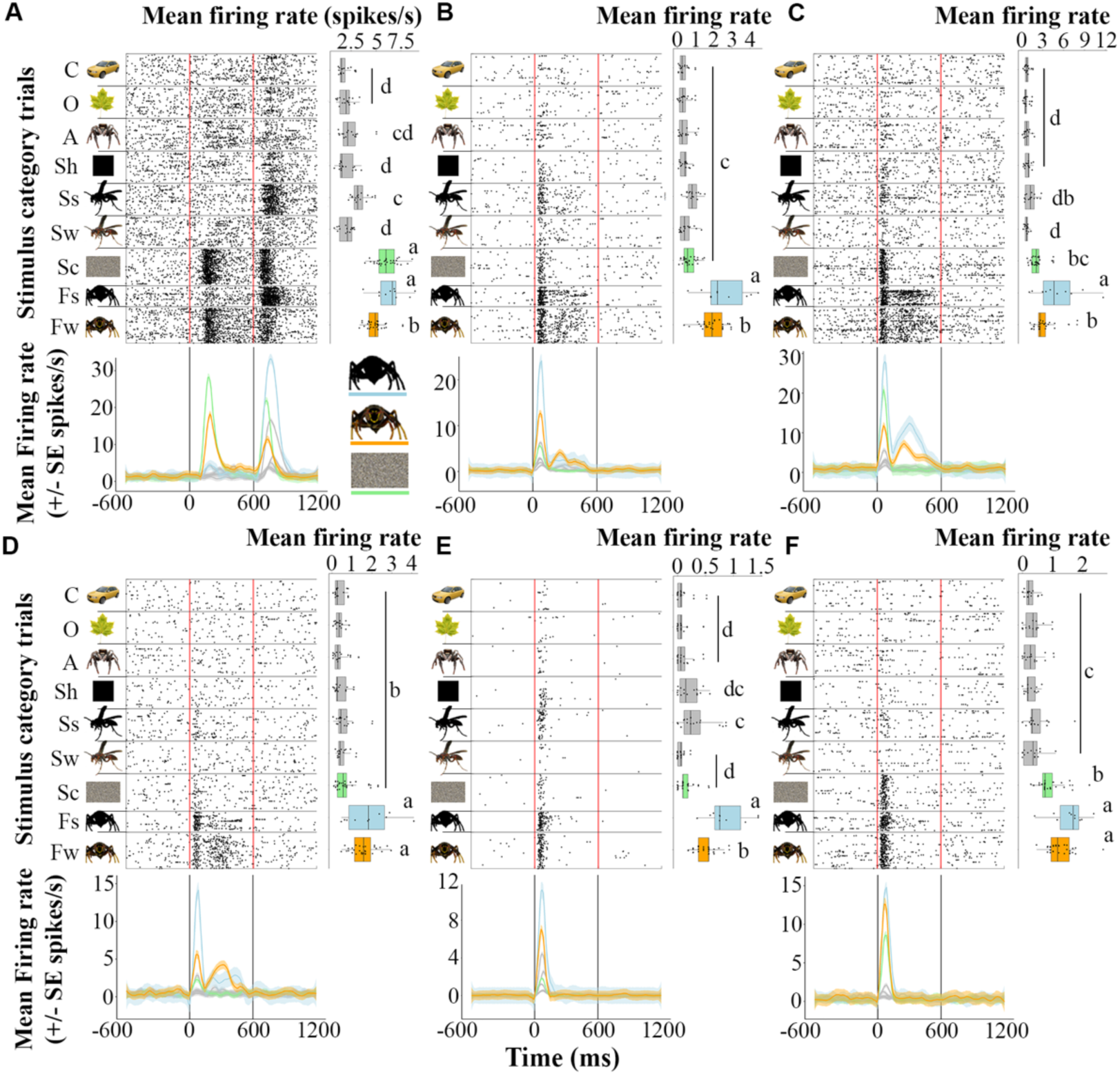
Additional stimulus set 1 example front-facing wasp selective units. Additional example raster plots organized the same as Fig. 2C-E and shown stimulus set 1; showing all 2296 total object presentations for the 706 unique object stimuli sorted by the 9 stimulus categories highlighted on the y-axis. Stats represent ANOVA analysis of mean firing rate ∼ stimulus category and letters denote p<0.05 significant difference by Tukey HSD post hoc analysis. Color denotes stimulus category for each unit’s response front-facing wasp is highlighted in orange, front-facing wasp silhouette in blue, and scramble in green. (**A**) Example *wasp cell* unit 186 from wasp 4. (**B**) Example *wasp cell* unit 330 from wasp 3 with persistent firing to wasp shape. (**C**) Example *wasp cell* unit 366 from wasp 3 with persistent firing to wasp shape. (**D**) Example *wasp cell* unit 151 from wasp 3 with persistent firing to wasp shape. (**E**) Example *wasp cell* unit 329 from wasp 3 with phasic firing to wasp shape. (**F**) Example *wasp cell* unit 360 from wasp 3 with phasic firing to wasp shape and color. Acronyms: C=car, O=other, A=non-wasp arthropod, Sh=shape, Ss=side wasp silhouette, Sw=side wasp, Sc=scramble, Fs=front-facing wasp silhouette, Fw=front-facing wasp.

**Fig. S2.**
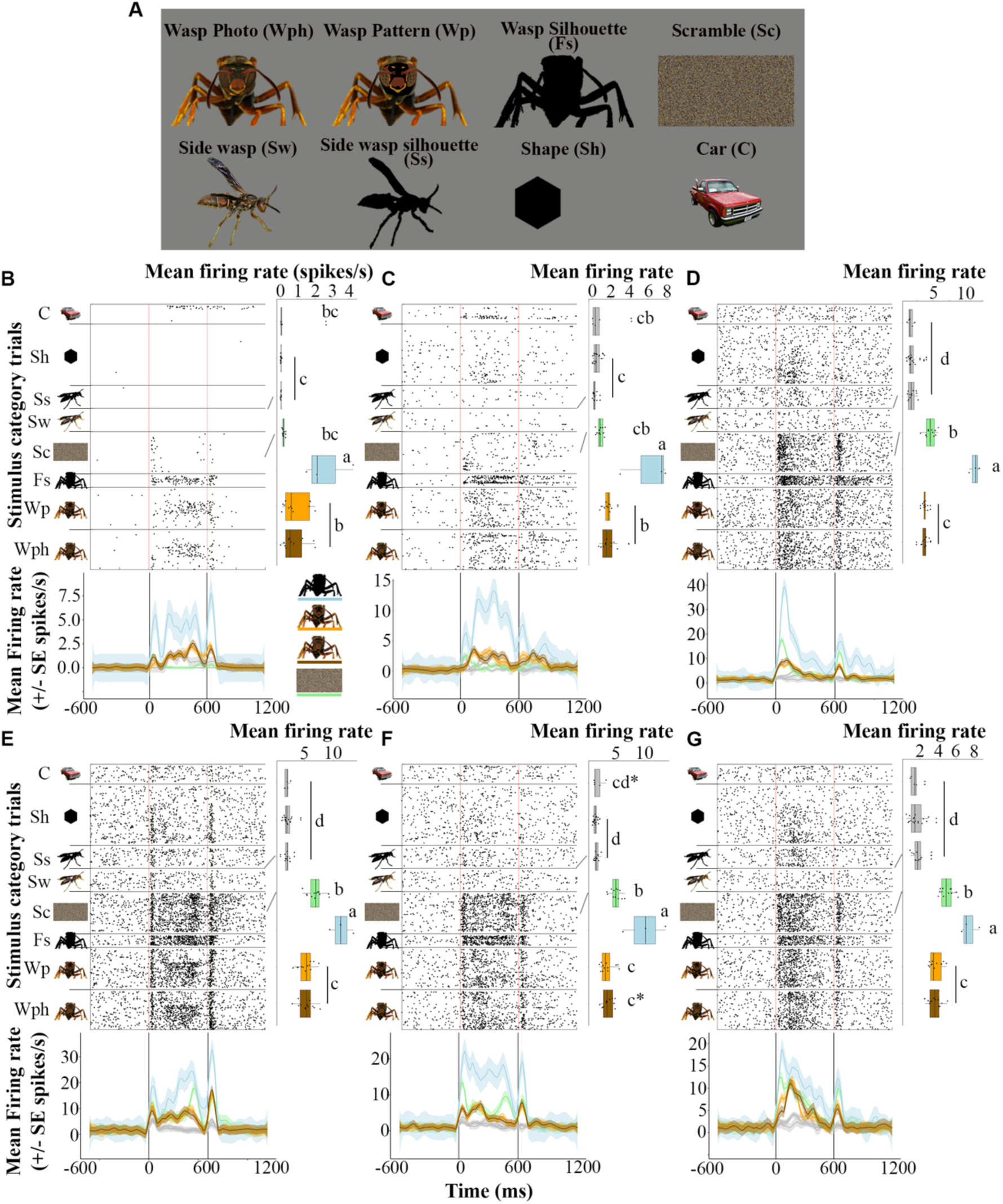
Stimulus set 2 and example front-facing wasp selective units. (**A**) Example images of 8 primary stimulus categories used in stimulus set 2. (**B**-**G**) Example raster plots above with persistent firing to wasp shape and color from wasp 5, organized the same as Fig2C-E but for units shown stimulus set 2. Each row in the raster plot is a unique stimulus presentation showing 1184 of the 1664 total presentations, made up of 74 of the total 104 unique stimuli sorted by the 8 stimulus categories highlighted on the y-axis (omitted antennaless and solid colored face stimuli). Stats represent ANOVA analysis of mean firing rate ∼ stimulus category and letters denote p<0.05 significant difference by Tukey HSD post hoc analysis. Color denotes stimulus category for each unit’s response to front-facing wasp with standardized face pattern highlighted in orange, front-facing wasp face photo highlighted in brown, front-facing wasp silhouette in blue, and scramble in green. (**B**) Example *wasp cell* unit 286. (**C**) Example *wasp cell* unit 84. (**D**) Example *wasp cell* unit 287. (**E**) Example *wasp cell* unit 341. (**F**) Example *wasp cell* unit 266. (**G**) Example *wasp cell* unit 264. Acronyms: C=car, Sh=shape, Ss=side wasp silhouette, Sw=side wasp, Sc=scramble, Fs=front-facing wasp silhouette, Wp=front-facing wasp with cartoonized pattern, Wph=front-facing wasp photo.

**Fig. S3.**
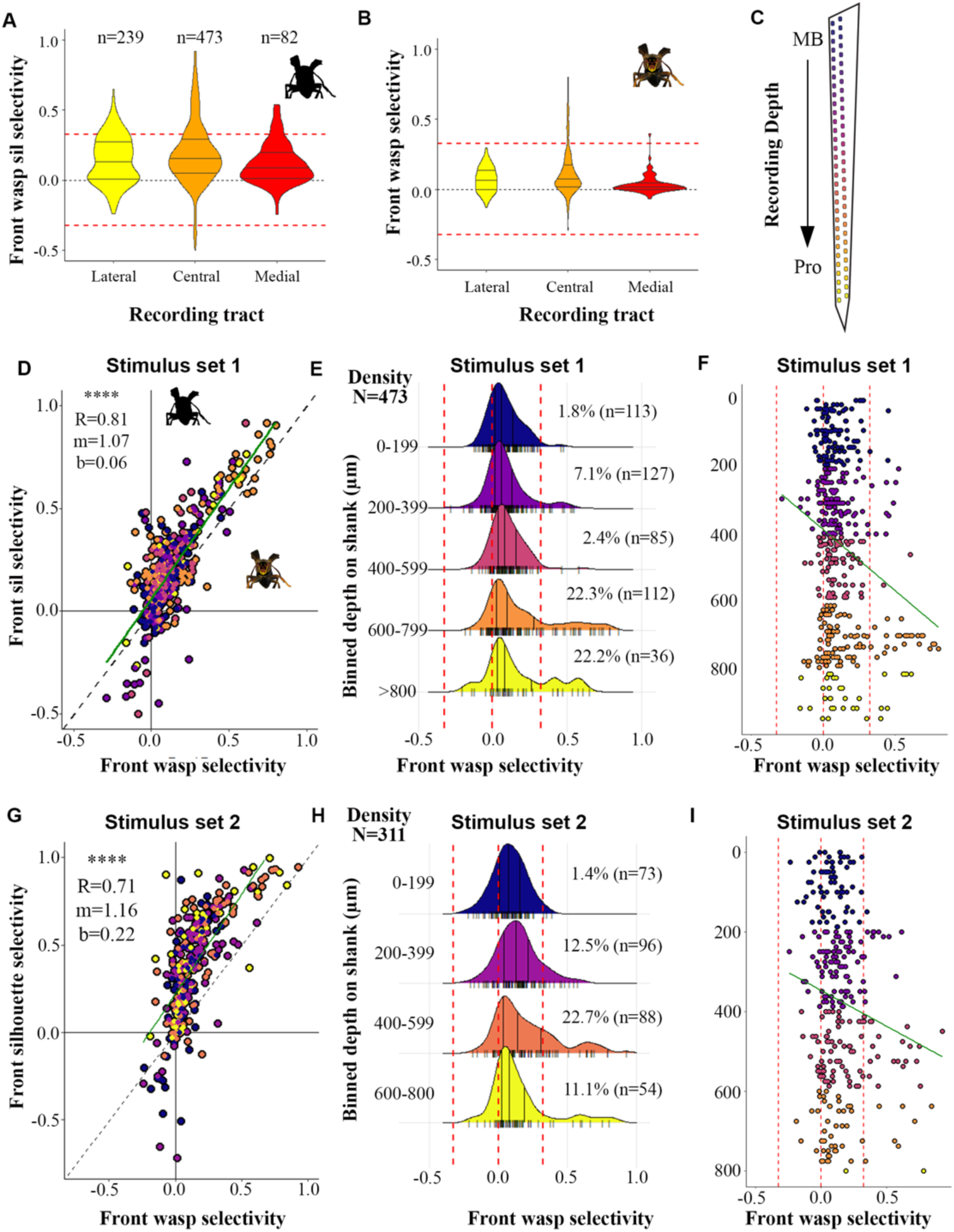
Relationship by depth and consistent relationship of front-facing wasp shape correlation across stimulus sets. (**A**) Distribution of selectivity to front-facing wasp silhouette images across the three recording tracts. Number above each violin plot denotes the number of visually responsive units recorded in each tract. See Fig. 1D for recording tract schematic. (**B**) Distribution of selectivity to front-facing wasp images across the three recording tracts. (**C**) Enlarged schematic of the silicon probe shank within a dashed box, channel depth is color coded for each recording channel location with deeper more warmer colors denoting lateral protocerebrum colors and cooler colors denoting mushroom body recording cites. (**D**) Relationship of front-facing wasp silhouette selectivity and front-facing wasp selectivity across 473 units recorded in the central tract. Stats denote Pearson correlation analysis and linear fit. Colors denote recording location binned by depth along the recording shank. (**E**) Distribution of selectivity to front-facing wasp stimuli in 200 μm depth bins for units recorded in the central tract. Colors denote depth as displayed in C and D. Numbers denote the number of visually responsive units recorded at each depth and percentages denote the percentage of units that show >=2x selectivity to front-facing wasp stimulus category. (**F**) Front-facing wasp selectivity for all units recorded from the central tract presented stimulus set 1 and the depth along the recording shank on which they were recorded instead of binned as in E. Green line denotes linear fit model showing significantly increased selectivity to front-facing wasp images as you move along recording depth. (**G**) Relationship of front-facing wasp silhouette selectivity and front-facing wasp selectivity across 311 units recorded in the central tract shown stimulus set 2. Stats denote Pearson correlation analysis and linear fit. Colors denote recording location by depth along the recording shank. (**H**) Distribution of selectivity to front-facing wasp stimuli in 200 μm depth bins for units recorded in the central tract show stimulus set 2. Colors denote depth as displayed in C and G. Numbers denote the number of visually responsive units recorded at each depth and percentages denote the percentage of units that show >=2x selectivity to front-facing wasp stimulus category. (**I**) Front-facing wasp selectivity for all units recorded from the central tract presented stimulus set 2 and the depth along the recording shank on which they were recorded instead of binned as in H. Green line denotes linear fit model showing significantly increased selectivity to front-facing wasp images as you move deeper along recording tract. Acronyms: MB= mushroom body calyx, Pro=protocerebrum.

**Fig. S4.**
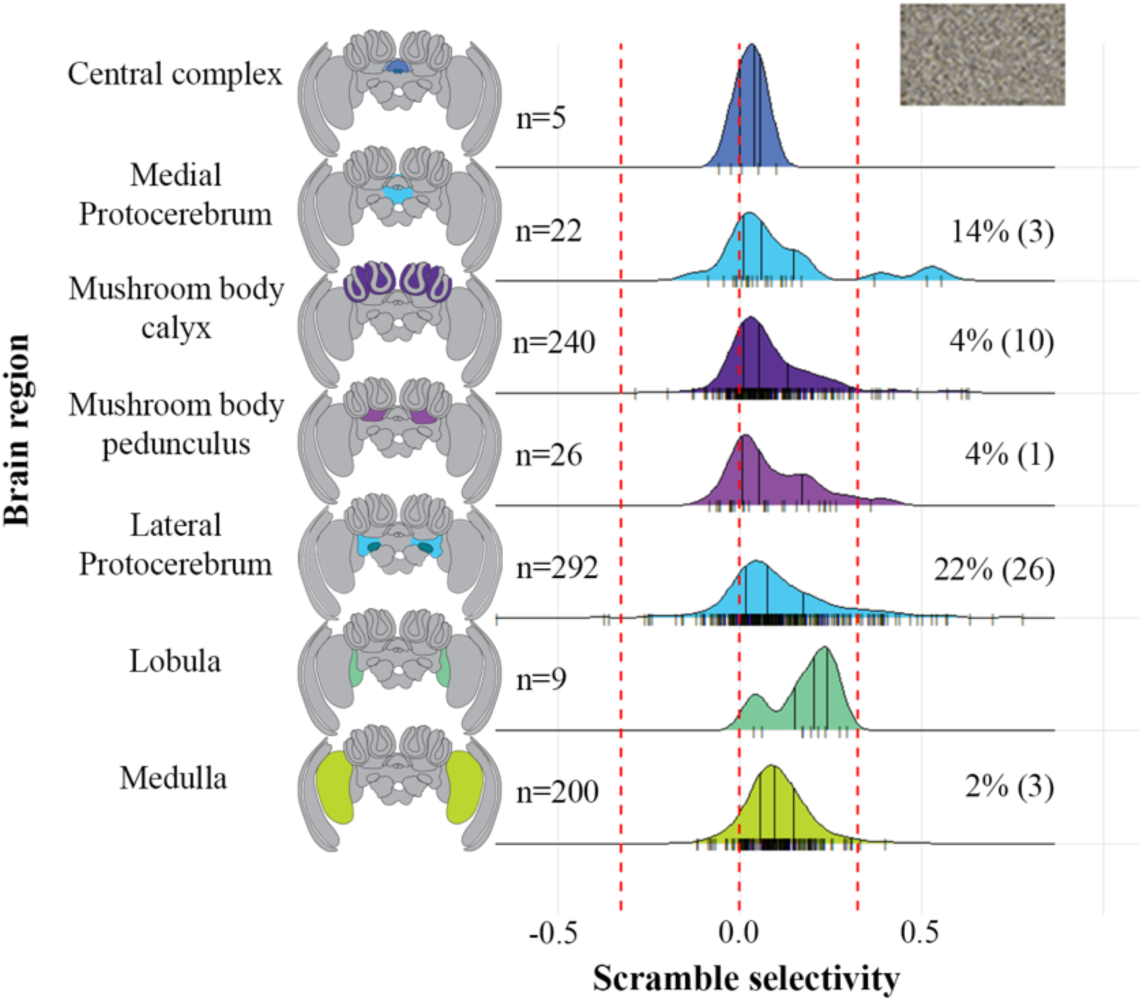
Distribution of selectivity to scramble images across regions of the wasp brain. Numbers to the right denote the total number of visual units record in each neuropil and percentages (and number of units in parentheses) to the right denote the percentage of units in each neuropil that show >=2x selectivity to scramble stimuli.

**Fig. S5.**
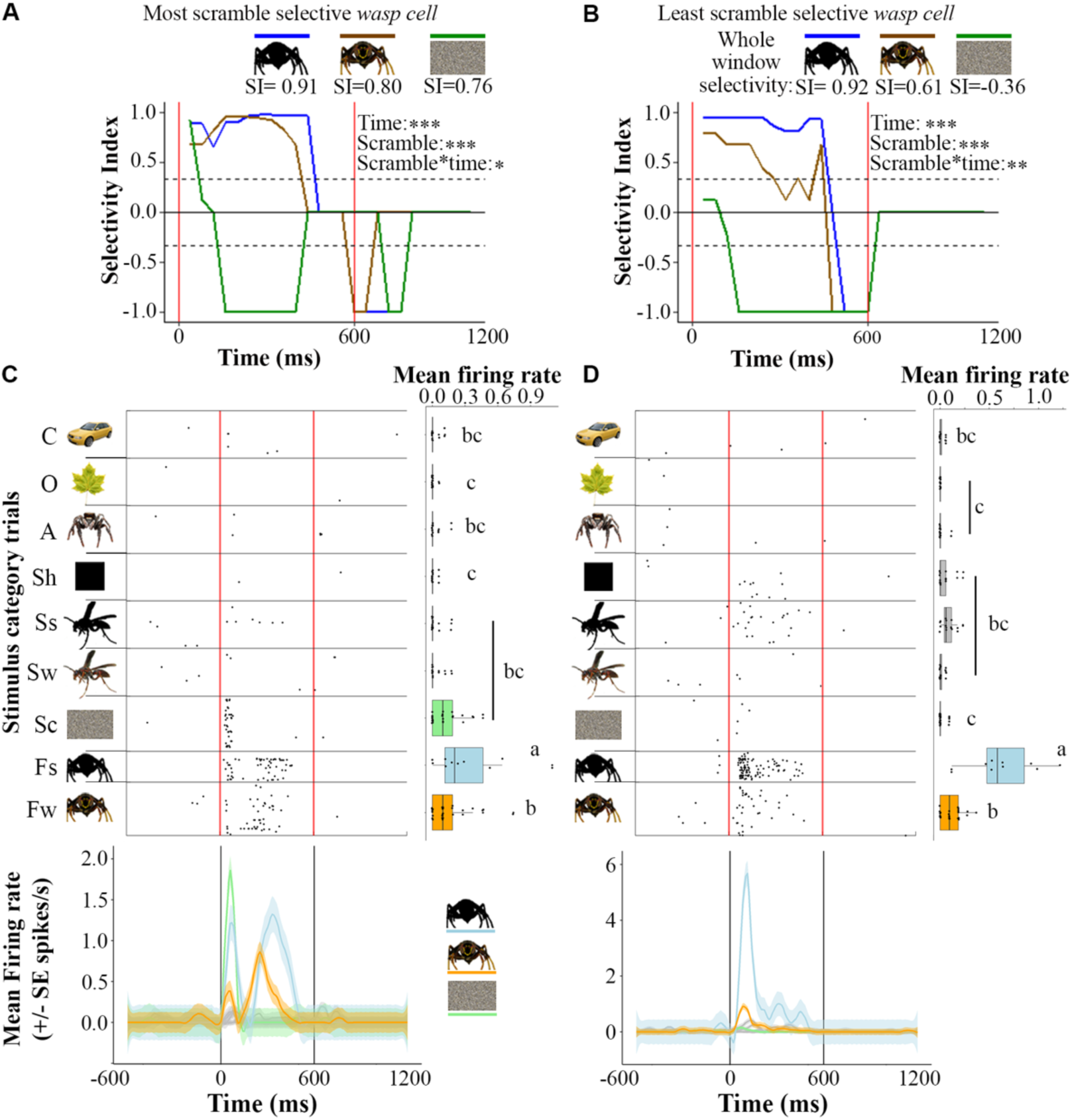
Example *Wasp cell* selectivity over time for the most and least scramble selective *wasp cells*. (**A**) Example *wasp cell* 154 from wasp 3 whose raster is plotted in C and its selectivity in 40ms bin windows over the whole stimulus response window (0-1080 ms) to front-facing silhouette (blue), wasp (orange), and scramble (green). Selectivity index (SI) values for this cell’s whole response window are displayed above. Stats denote significance running GLM analysis, see supplementary methods for additional details. (**B**) Example *wasp cell* 281 from wasp 3 whose raster is plotted in D and its selectivity in 40 ms bin windows as in A. Line plots in A and B are smoothed using a median selecting a value across 3 bins. (**C**-**D**) Example raster plots above, organized the same as Fig. S1. (**C**) Example *wasp cell* unit 154 from wasp 3, which is the most scramble selective among measure *wasp cells*. (**D**) Example *wasp cell* unit 281 from wasp 3, which is the least scramble selective among measure *wasp cells*.

**Fig. S6.**
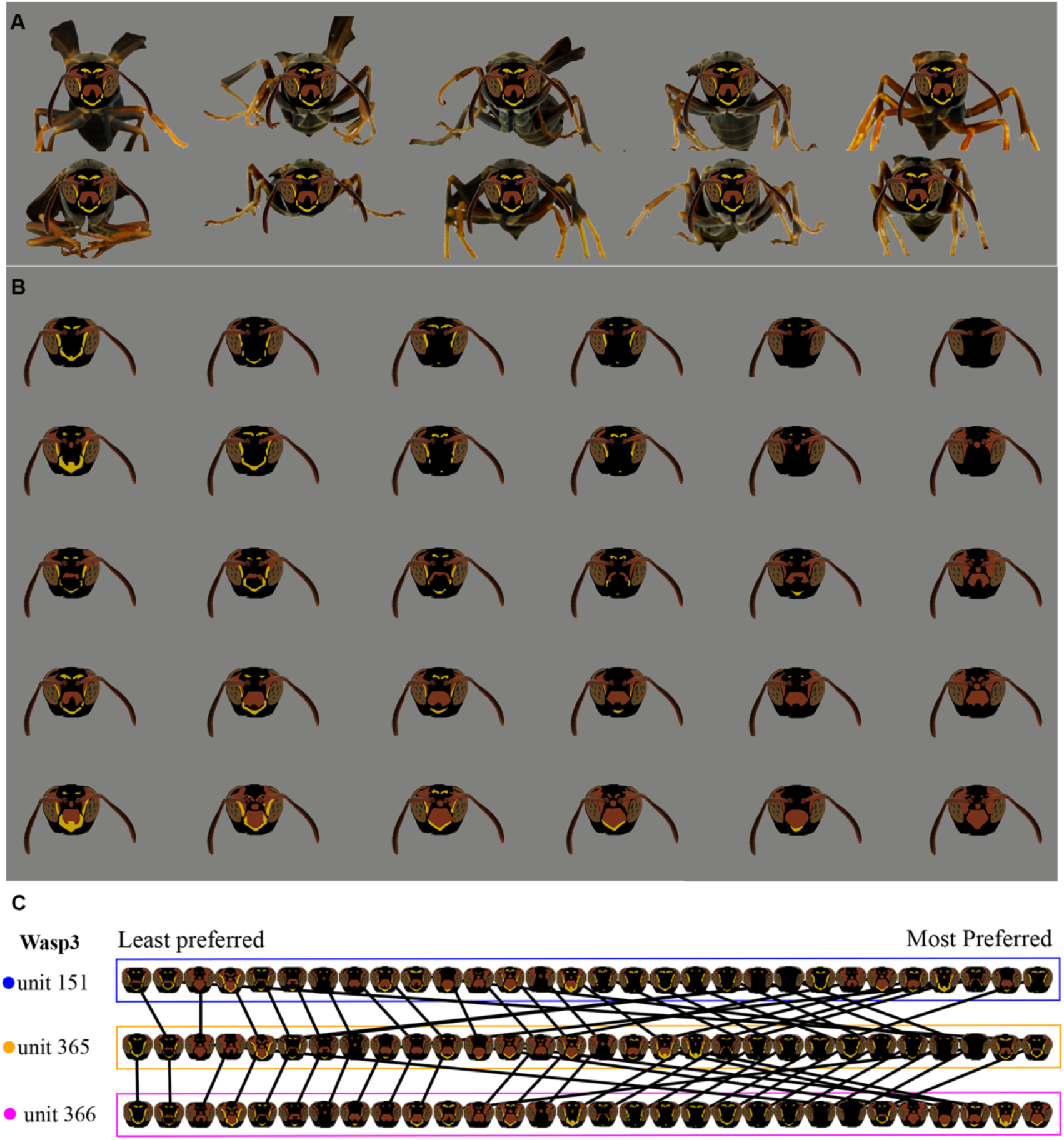
Unique wasp face and body stimuli used in stimulus set 1 and unique facial tuning of example *wasp cells*. (**A**) Example patterned front-facing wasp stimuli images. Displayed are the same wasp facial pattern presented on each of the 10 unique shown body postures. (**B**) Example of the 30 unique wasp facial patterns presented in stimulus set 1, note bodies removed for clarity but stimuli were each presented on 10 unique bodies. (**C**) Three example *wasp cells* recorded from wasp 4 which significantly vary in their firing across faces, raster plots for each cell can be found in Fig. 2E and fig. S1C & D. Each row sorts faces from least fired to most fired from left to right among the 30 presented wasp faces stimuli. Lines connect the same faces across units. The tangle of lines suggests non-uniform preferences across units.

**Fig. S7.**
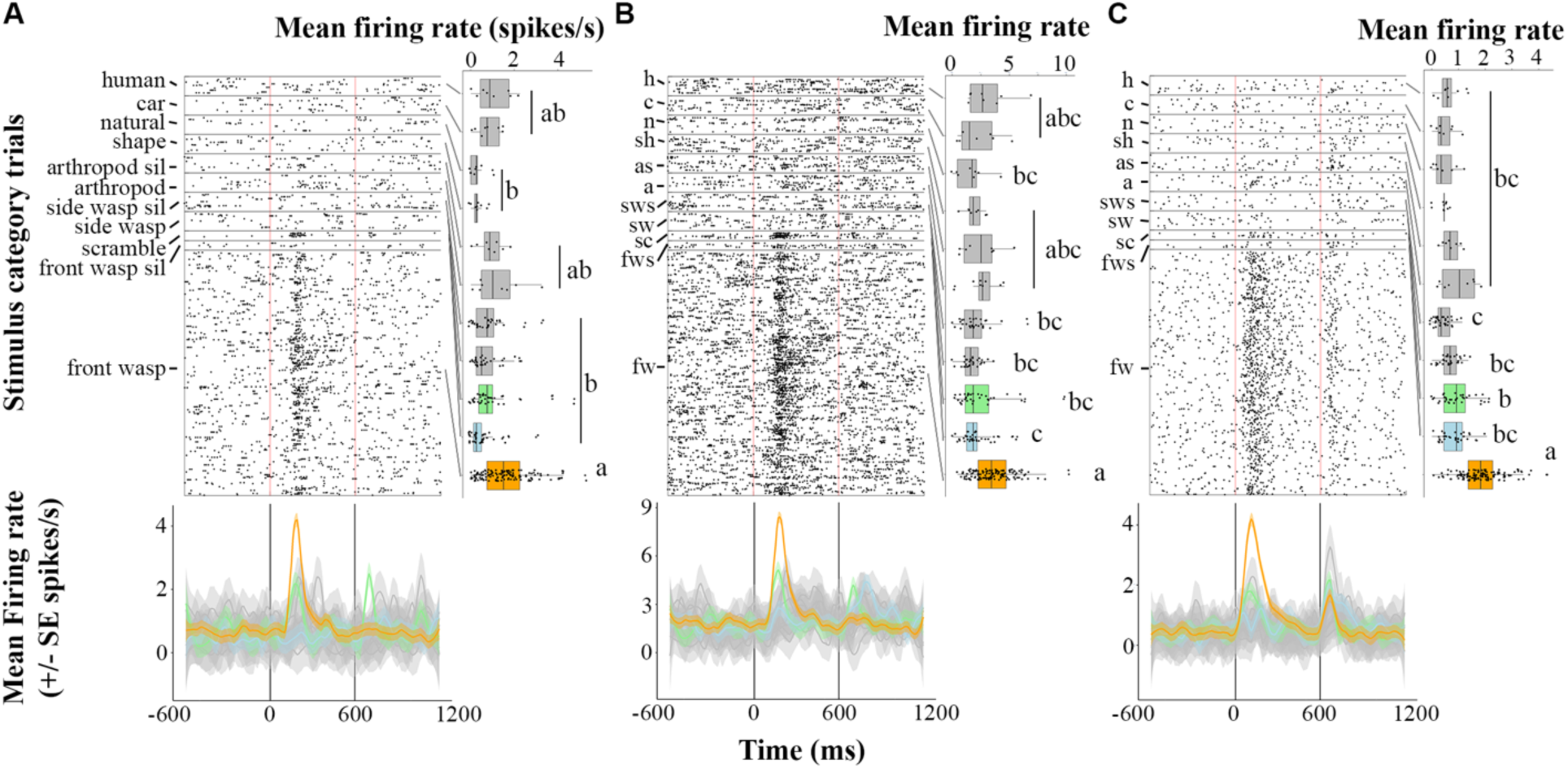
Example raster plots for example stimulus set 3 *wasp cells* shown in Fig 5. (**A**-**C**) Example *wasp cell* raster plots shown stimulus set 3 and found in Fig. 5B-M, organized the same as Fig 2C-E. Each row in the raster plot is a unique stimulus presentation showing 1960 of the 3400 total presentations, made up of 196 of the total 340 unique object stimuli sorted by the 11 stimulus categories highlighted on the y-axis. Stats represent ANOVA analysis of mean peak firing rate ∼ stimulus category and letters denote p<0.05 significant difference by Tukey HSD post hoc analysis. Color denotes stimulus category for each unit’s response, front-facing wasp in orange, front-facing wasp silhouette in blue, and scramble in green. (**B**) Example *wasp cell,* unit 51 from wasp 6. (**C**) Example *wasp cell,* unit 254 from wasp 7. (**D**) Example *wasp cell,* unit 365 from wasp 7.

**Fig. S8.**
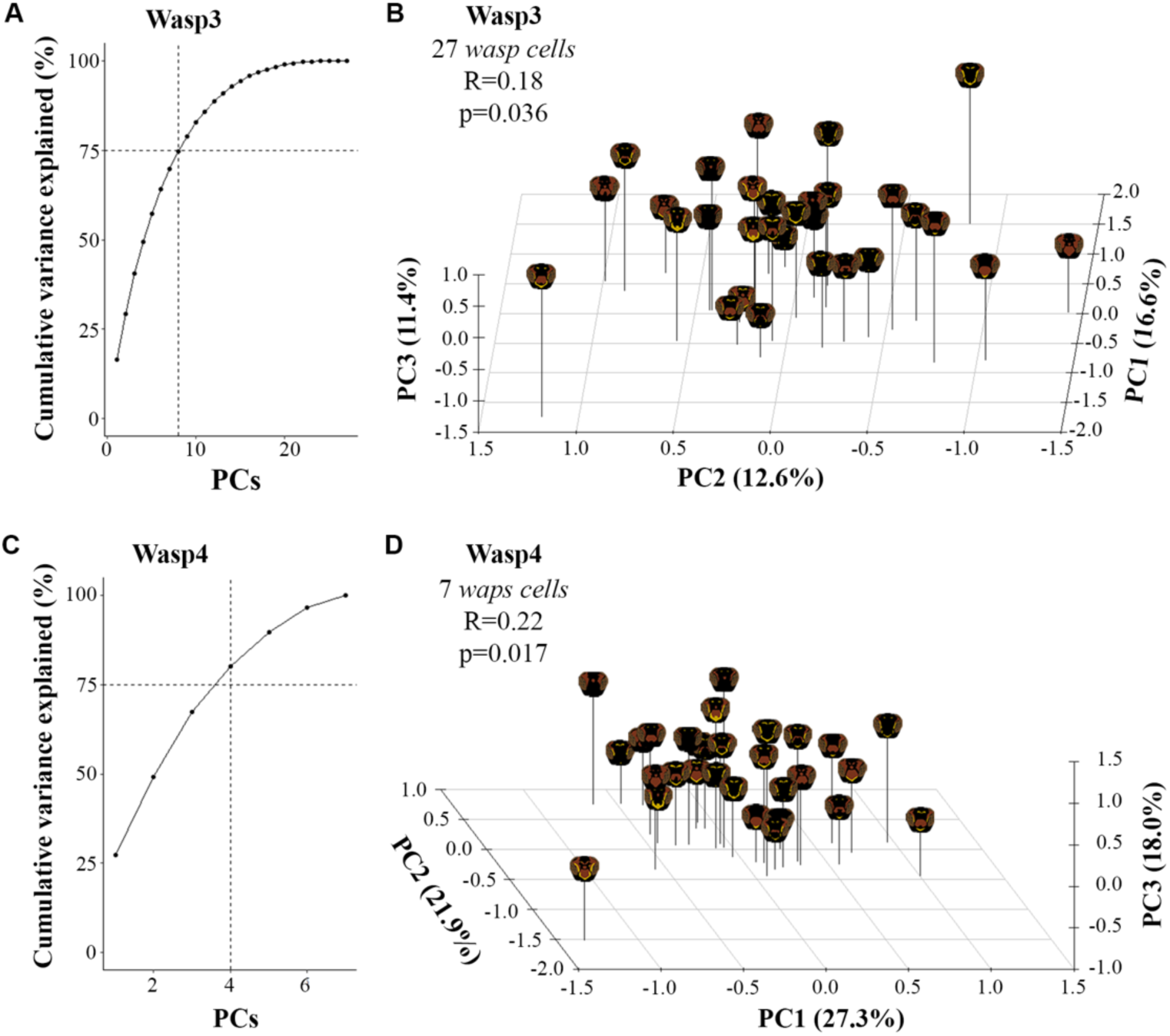
Additional single animal *wasp cell* PCA analysis plots matching Fig. 5M & N. (**A**) Plot of the cumulative variance explained for each principal component (PC) of the principal component analysis (PCA) for *wasp cells* from wasp 3. (**B**) Plot of face locations for the first 3 PCs of the PCA for the 27 *wasp cells* from wasp 3 using their z-score peak firing to the 30 wasp faces shown in stimulus set 1. (**C**) Plot of the cumulative variance explained for each PC of the PCA for *wasp cells* from wasp 4. (**D**) Plot of face locations for the first 3PCs of the PCA for the 7 *wasp cells* from wasp 4 using their z-score peak firing to the 30 wasp faces shown in stimulus set 1. Lines in A & C denote 8 and 4 PCs respectively being required to explain 75% of the total variance present across the 27 and 7 *wasp cells* respectively. Stats in B and D denote significant correlation of facial distances across neural firing PC space and facial phenotype PC space via Mantel test with 100,000 permutations.

